# Upregulation of the AcrAB_2_NodT efflux pump confers antibiotic resistance at the cost of collateral metal sensitivity

**DOI:** 10.1101/2025.07.17.665315

**Authors:** Marine Ote, Lorine Lardinois, Erika Hendrickx, Marc Dieu, Dukas Jurėnas, Jean-Yves Matroule

**Affiliations:** Research Unit in Biology of Microorganisms (URBM), Department of Biology, Namur Research Institute for Life Sciences (NARILIS), University of Namur, Namur, Belgium; Center for Microscopy and Molecular Imaging (CMMI), Université Libre de Bruxelles, Gosselies, Belgium; MaSUN, Mass Spectrometry Facility, University of Namur, Namur, Belgium; Bacterial Genetics and Physiology, Faculté des Sciences, Université Libre de Bruxelles (ULB), Gosselies, Belgium. WEL Research Institute, Wavre, Belgium

**Keywords:** Antibiotics, antibiotic resistance, heavy metals, metal sensitivity, copper, stress response, efflux pumps, *Caulobacter*

## Abstract

Bacterial resistance to antibiotics (AB) such as β-lactams, fluoroquinolones, and aminoglycosides often emerges through mutations that alter AB targets, reduce membrane permeability, or increase the activity of AB-modifying enzymes and efflux pumps. Yet the physiological costs associated with AB resistance remain poorly understood. In *Caulobacter vibrioides*, a Δ*tipR* mutant that which constitutively upregulates the RND pump AcrAB_2_nodT, displays heightened sensitivity to copper (Cu), revealing a physiological vulnerability driven by the energetic burden imposed by excessive efflux activity. Deletion of *acrAB_2_nodT* in the Δ*tipR* background restored Cu resistance to wild-type levels, confirming the role of pump overexpression in metal sensitivity. Morphological and microscopy analyses revealed that pump overexpression leads to cell envelope defects and compromised fitness. To disentangle the effects of pump abundance from efflux activity, we engineered an AcrB_2_ transport-impaired mutant. This variant also rescued Cu resistance, demonstrating that high expression and active transport contribute to the observed toxicity. Notably, this sensitivity was not limited to Cu; the Δ*tipR* mutant also exhibited increased susceptibility to other transition metals, including zinc (Zn), nickel (Ni) and cadmium (Cd), suggesting a broader vulnerability linked to metal stress. Mechanistically, pump overexpression depleted the proton motive force, reduced ATP levels, and impaired motility, all of which are essential for Cu stress adaptation. This physiological tradeoff highlights the importance of precise efflux regulation and reveals a potential therapeutic vulnerability: targeting the cost of pump upregulation could enhance the efficacy of antimicrobial treatments.

**Importance:** While efflux pumps primarily protect bacteria from AB, their excessive activity can impose a fitness cost. This study underscores a crucial tradeoff between resistance and cellular fitness by connecting Cu sensitivity to heightened AcrAB_2_NodT efflux pump expression in *Caulobacter vibrioides*. This work challenges the assumption that more efflux always benefits the cell and shows that unregulated pump activity can undermine survival under metal stress. These findings broaden our understanding of bacterial stress physiology and suggest new ways to combat antimicrobial resistance by targeting the hidden costs of resistance mechanisms. Understanding this balance between resistance and fitness opens new perspectives for combination therapy against multidrug-resistant bacteria.

## Introduction

Antimicrobial resistance (AMR) represents an urgent and growing global health threat, with projections estimating up to 8.22 million AMR-associated deaths annually by 2050 (1). Although AMR is well-studied in clinical settings, environmental contamination is now recognized as a major driver of the spread and evolution of antimicrobial resistance. Human activities, including insufficiently treated wastewater discharge, the extensive use of AB in agriculture, and various industrial practices, have led to the accumulation of AB in natural habitats (2). Consequently, environmental bacteria are frequently exposed to other pollutants, such as heavy metals, organic toxins, and industrial chemicals. This chronic exposure exerts selective pressure that promotes the development and dissemination of diverse resistance mechanisms (3). Among these, Resistance-Nodulation-Division (RND) efflux pumps have emerged as key elements of bacterial defense against toxic compounds (4). RND systems, prevalent in Gram-negative bacteria, are tripartite complexes spanning the inner and outer membranes and are powered by the proton motive force (PMF). These complexes consist of an inner membrane RND transporter, a periplasmic membrane fusion protein (MFP), and an outer membrane factor (OMF), functioning together to export a broad range of harmful substrates out of the cell (5).

RND efflux pumps are generally classified into two major subfamilies based on substrate specificity: the hydrophobic and amphiphilic efflux RND (HAE-RND) pumps, which primarily expel AB and detergents, and the heavy metal efflux RND (HME-RND) pumps, which mediate metal ion resistance (6). In the model organism *E. coli*, the HAE-RND pump AcrAB-TolC and the HME-RND pump CusCFBA have been extensively characterized for their respective roles in AB and heavy metal detoxification, respectively (7, 8). These systems are tightly regulated at the transcriptional level, typically via local repressors such as AcrR for AcrAB-TolC and CusR for CusCFBA (7, 8). Notably, clinical isolates often harbor mutations in these regulatory elements, leading to constitutive overexpression of efflux systems and multidrug-resistant phenotypes (9).

While RND efflux pumps are extensively studied for their role in antimicrobial resistance, particularly in clinical pathogens, the broader physiological consequences of their overexpression remain poorly understood (10). This gap is especially pronounced in environmental bacteria, where the regulation of efflux systems under complex and fluctuating stressors has received little attention (11). As a result, the potential trade-offs between resistance and bacterial fitness in natural ecosystems remain largely unexplored.

In this study, we investigate the fitness consequences of unregulated expression of the AcrAB_2_NodT efflux pump in the environmental bacterium *C. vibrioides*. Our results reveal that the upregulation of this HAE-RND pump induces metal sensitivity, accompanied by morphological defects, envelope remodeling, and metabolic reprogramming driven by membrane depolarization and ATP depletion. These findings underscore the importance of tightly regulated efflux activity and highlight a previously underappreciated trade-off in resistance evolution. Moreover, this work suggests that dual-targeting strategies involving AB and metals may offer a promising alternative for future antimicrobial therapies.

## Results

### Collateral Cu sensitivity of the Δ*tipR* mutant

A genetic screen performed on a *C. vibrioides* mini-Tn5 transposon mutant library to seek Cu-sensitive mutants led to the identification of the *tipR* gene (*CCNA_00852*) (Fig. 1A). TipR encodes a one-component transcriptional regulator that was shown to repress the expression of the neighbouring efflux pump AcrAB_2_NodT (12). To validate the genetic screen, we deleted the *tipR* gene in the WT strain. We monitored the growth of the resulting Δ*tipR* mutant (hereafter referred to as Δ*R*) over 24 h in rich liquid PYE medium under moderate (150 µM CuSO₄), sub-high (175 µM CuSO₄), and high (190 µM CuSO₄) Cu stress conditions and on solid PYE medium under moderate (100 µM CuSO₄) Cu stress. The Δ*R* mutant exhibited an increased sensitivity to moderate Cu stress in solid (Fig. 1B) and liquid (Fig. 1C) PYE medium but displayed a WT-like growth under control condition (Fig. S1A). Cu sensitivity correlated with increasing CuSO₄ concentrations, with the Δ*R* mutant displaying reduced fitness at high Cu stress (Fig. S1A). Ectopic expression of *tipR* from the low-copy plasmid pMR10 under the control of the constitutive *lac* promoter (Δ*R+R*) restored the WT phenotype (Fig. 1B-C), indicating that *tipR* deletion is not associated with polar effects.

**Figure 1.**
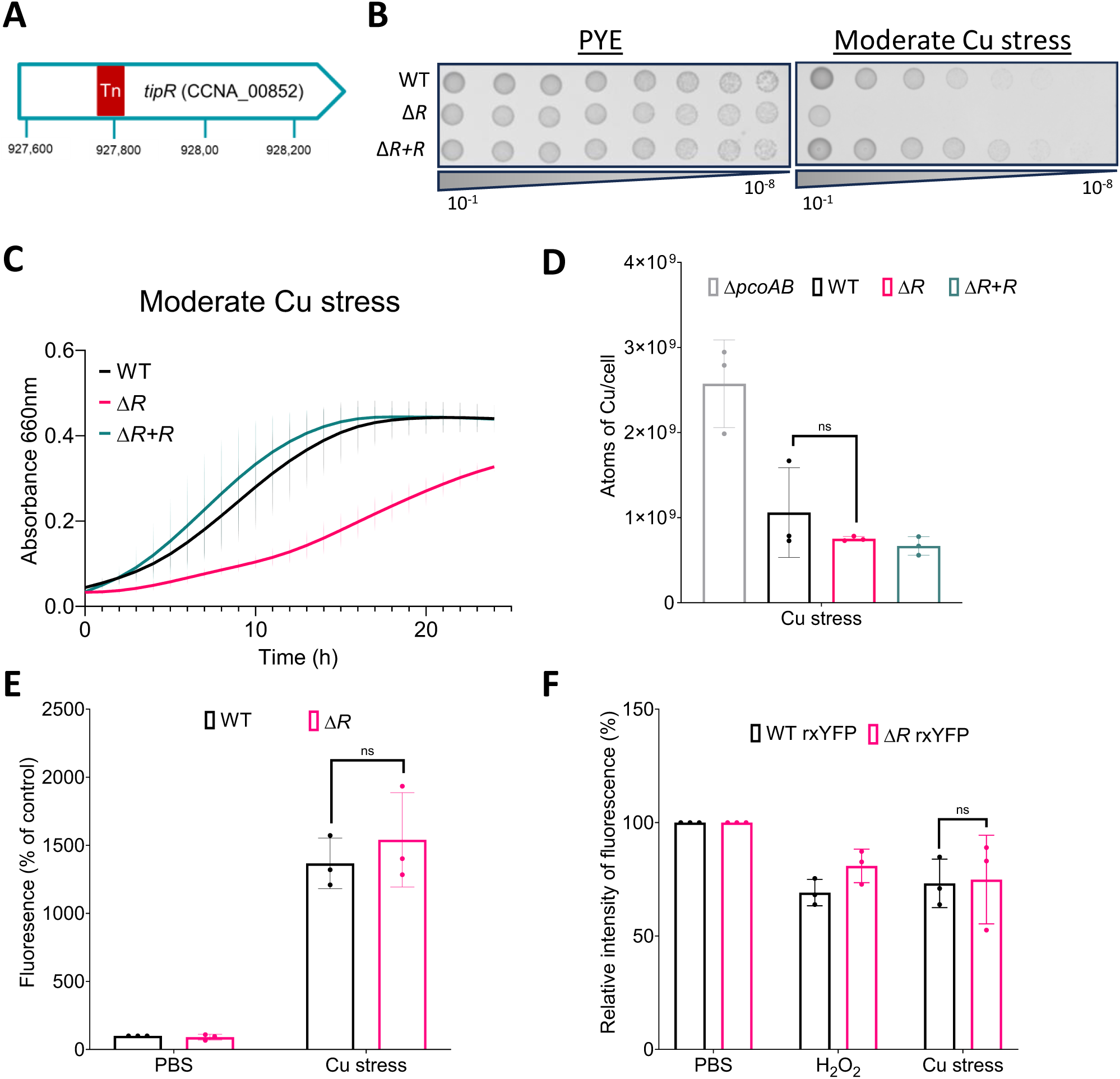
Cu sensitivity of the Δ*R* mutant is independent of Cu accumulation and ROS production. **A.** Schematic of the *tipR* gene with insertion of a Mini Tn-5 found during a genetic screen for Cu sensitivity. **B.** Viability assay on PYE plates of WT, Δ*R*, and Δ*R+R* strains, under control and 100 µM CuSO_4_ conditions. **C.** Growth profiles at an OD_660_ nm of WT, Δ*R*, and Δ*R+R* strains in PYE medium supplemented with 150 µM CuSO_4_. **D.** Number of Cu atoms per cell exposed to 175 µM CuSO_4_ excess for 1 h. **E**. Relative fluorescence intensity of WT, Δ*R*, and Δ*R+R* strains incubated with 5 μM CellROX deep red probe and exposed to 175 μM CuSO_4_. **F.** Relative fluorescence intensity of WT, Δ*R*, and Δ*R+R* strains expressing the rxYFP biosensor and exposed to 175 μM Cu or 400 μM H_2_O_2_. All data are represented as the mean ± SD, with at least three biological replicates. *p* values were calculated using ANOVA combined with Sidak’s, or Dunnett’s multiple comparison test (\*\*\*\**p* < 0.0001, \*\*\**p* < 0.001, \*\**p* < 0.01, \**p* < 0.05).

To determine whether the sensitivity of the Δ*R* mutant was specific to Cu, we exposed the Δ*R* mutant to Zn, Ni and Cd. The Δ*R* mutant also exhibited an increased sensitivity to ZnSO₄ in liquid PYE medium (Fig. S1B), and showed enhanced sensitivity to ZnSO₄, NiSO₄, and CdSO₄ on solid medium (Fig. S1C). The rest of the study will be focusing on the Cu sensitivity.

We hypothesized that the Cu hypersensitivity of the Δ*R* mutant might result from increased intracellular Cu levels. To test this hypothesis, we quantified total intracellular Cu concentrations using atomic absorption spectroscopy (AAS). Surprisingly, the Δ*R* mutant exhibited no significant increase in Cu accumulation compared to the WT (Fig. 1D). Similarly, Cu levels in a mutant lacking the AcrAB_2_NodT efflux pump (Δ*ABT*) remained comparable to those of the WT (Fig. S1D). These results demonstrate that the Cu sensitivity of the Δ*R* mutant is neither due to elevated intracellular Cu nor to a direct role of AcrAB_2_NodT in Cu efflux, revealing an unexpected uncoupling between metal sensitivity and total cellular Cu content.

Owing to the ability of Cu to trigger ROS production (13), we decided to determine whether the increased Cu sensitivity of the Δ*R* mutant could result from a higher oxidative stress. Upon exposure to sub-high Cu stress, we observed an increase in fluorescence of the ROS-sensitive, turn-on probe CellROX deep red and the cytoplasmic redox-sensitive, turn-off biosensor rxYFP, confirming that Cu induces oxidative stress. However, no significant difference in CellROX (Fig. 1E) and rxYFP (Fig. 1F) fluorescence could be measured between the WT and Δ*R* strains, which is consistent with the similar Cu content of these two strains (Fig. 1D).

Together, these data indicate that the Cu sensitivity of the Δ*R* mutant is independent of increased Cu accumulation and oxidative stress.

### Inactivation of the AcrAB_2_NodT efflux pump restores Cu resistance in the Δ*tipR* mutant

To dissect the mechanisms underlying Cu toxicity in the Δ*R* mutant, we performed a suppressor screen in the Δ*R* mutant seeking mutants that would exhibit a WT-like Cu resistance. Nine suppressors were isolated with varying degrees of Cu resistance (Fig. S2A). Whole-genome sequencing identified four suppressor mutations as indels within *acrA*, which encodes the periplasmic component of the AcrAB_2_NodT efflux pump. The remaining five suppressor mutations were nucleotide substitutions located in the intergenic TipR-binding region between *tipR* gene and the *acrAB_2_nodT* operon (14) (Fig. 2A). TipR deletion triggers the upregulation of the AcrAB_2_NodT efflux pump (Fig. 2D) that confers resistance to penicillin (14) and cephalosporin (14), such as cefuroxime (CXM) (Fig. 2C). One could argue that the upregulation of the AcrAB_2_NodT efflux pump plays a key role in the collateral Cu sensitivity of the Δ*R* mutant.

**Figure 2.**
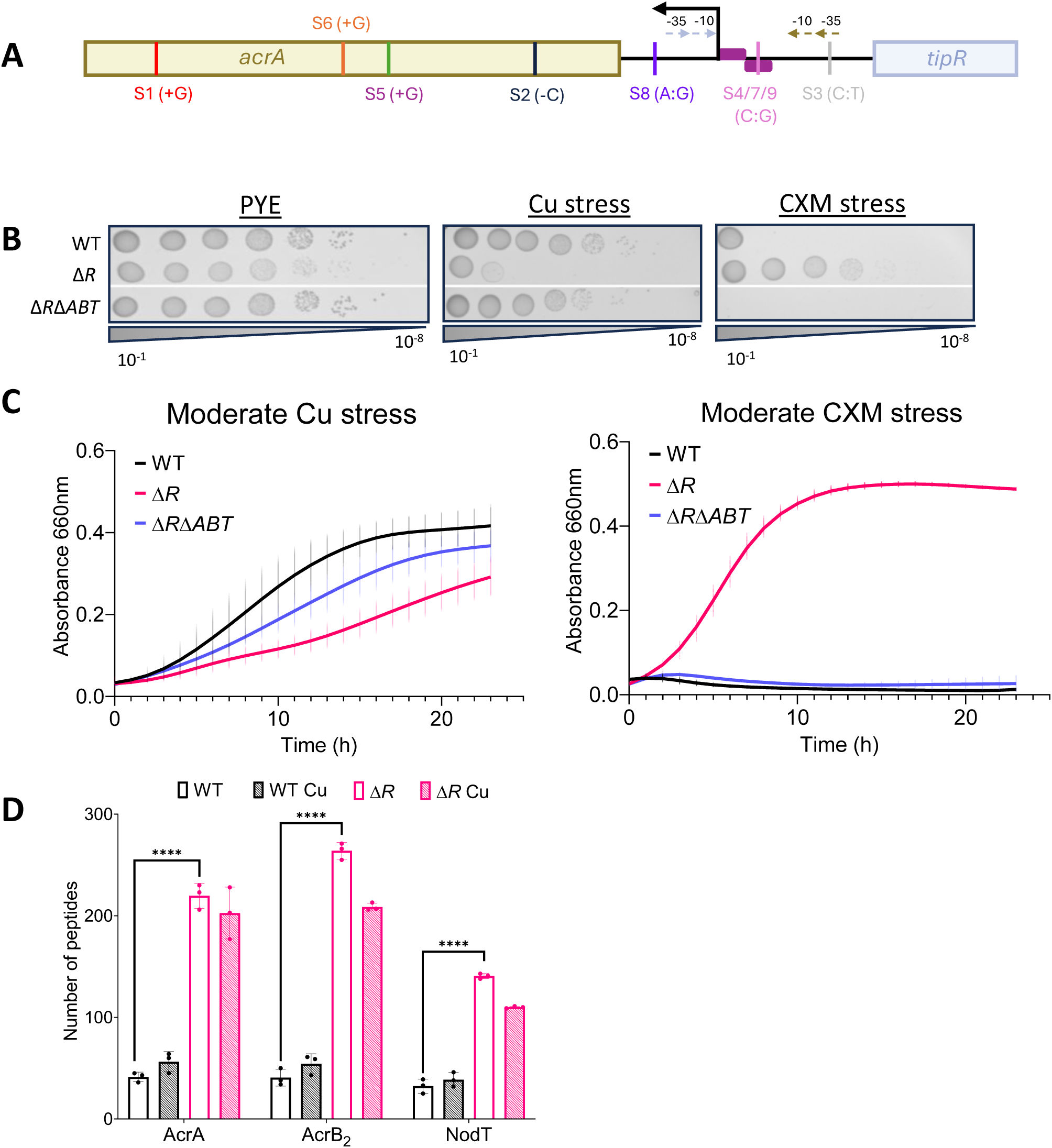
Inactivation of the AcrAB_2_NodT suppresses the Cu sensitivity of the Δ*R* mutant. **A.** Schematic of the *tipR* and *acrA* genomic region with the mutations found during a suppressor screen with 120 µM Cu. **B.** Viability assay on PYE plates of WT, Δ*R*, and Δ*R*Δ*ABT* strains, under control, 100 µM CuSO_4_ and 35 µg/ml CXM conditions. **C.** Growth profiles at an OD_660_ nm of WT, Δ*R*, and Δ*R*Δ*ABT* strains in PYE medium supplemented with 150 µM CuSO_4_ and 60 µg/ml CXM, respectively. **D.** Normalized total spectrum count of TipR and AcrAB_2_NodT protein in PYE medium, in combination with a 1 h exposure to 190 µM CuSO_4_. All data are represented as the mean ± SD, with at least three biological replicates. *p* values were calculated using ANOVA combined with Tukey’s multiple comparison test (\*\*\*\**p* < 0.0001, \*\*\**p* < 0.001, \*\**p* < 0.01, \**p* < 0.05).

Consistent with this hypothesis, a double *tipR*, *acrAB_2_nodT* operon (Δ*R*,*ABT*) mutant exhibits a WT-like Cu resistance in both solid (Fig. 2B) and liquid PYE medium (Fig. 2C), while losing its resistance to CXM (Fig. 2B–C). The abundance of AcrAB_2_NodT is not modulated by Cu excess though (Fig. 2D), which is consistent with the fact that this pump is not involved in Cu efflux (Fig. S1B).

### The upregulation of the AcrAB_2_NodT pump disrupts cell envelope integrity

The upregulation of the AcrAB_2_NodT efflux pump may cause a physiological burden in the bacterial envelope that would sensitize the cells to Cu. To test this hypothesis, we compared the cellular proteome of the WT and Δ*R* strains. Proteins involved in envelope remodeling are upregulated in the Δ*R* mutant, including fatty acid synthesis, proteases, transporters such as RND pumps, ABC transporters, TonB-dependent receptors and two-component systems (Fig. 3A). Consistent with this hypothesis, the Δ*R* mutant exhibits morphological defects under control and Cu stress conditions, including membrane blebbing, filamentation, and mislocalized stalks (Fig. S2B). These morphological defects were absent in the double Δ*R*Δ*ABT* mutant (Fig. S2B), indicating that they originate from AcrAB_2_NodT upregulation. The cell length and curvature were also reduced in the Δ*R* mutant under control condition (Fig. 3B) but were not further aggravated by Cu stress. Similar defects are observed when the WT strain is exposed to Cu stress (Fig. 3B).

**Figure 3.**
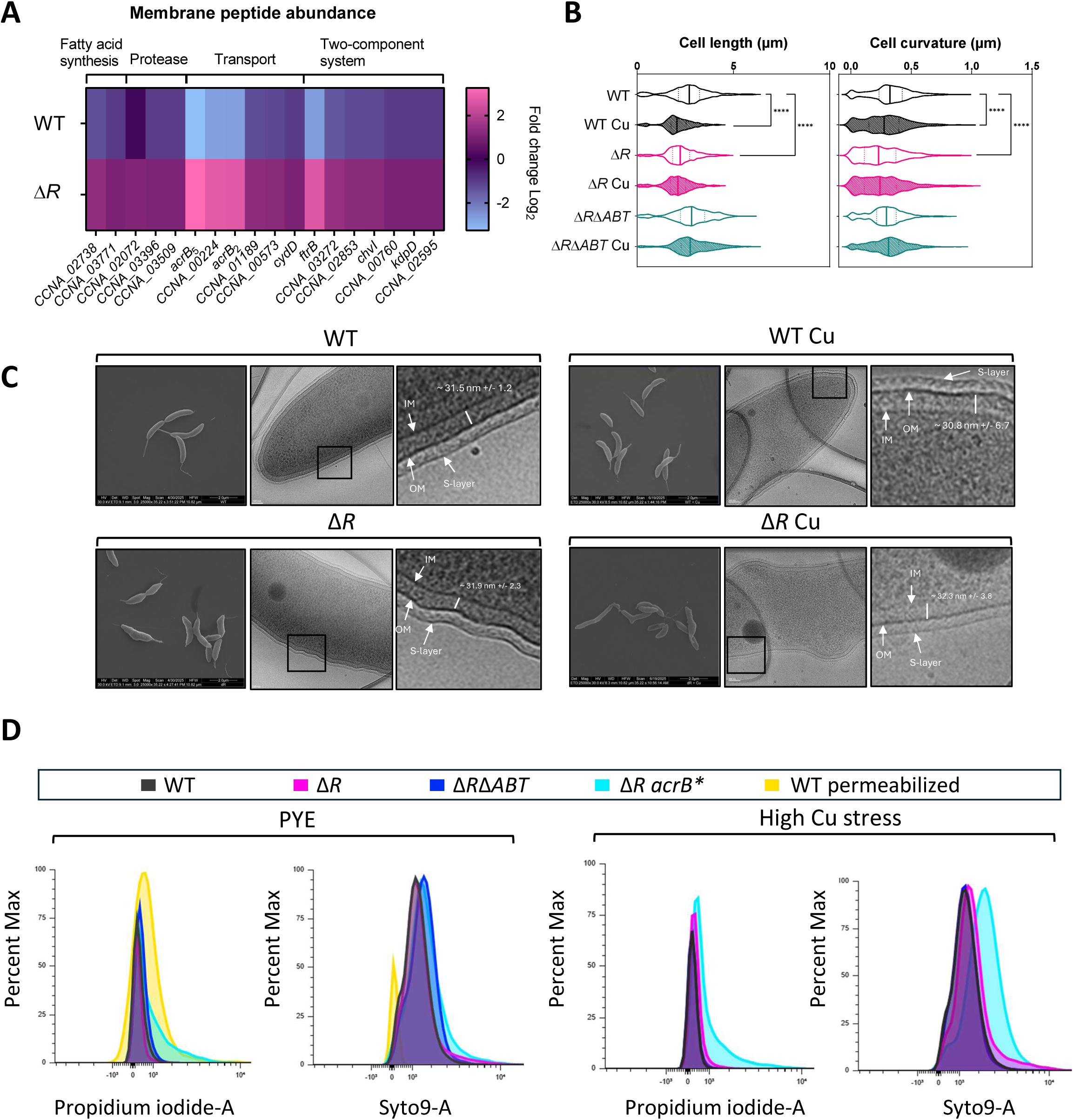
Upregulation of the AcrAB_2_NodT pump disrupts bacterial cell envelope integrity. **A.** Heat map depicting the fold change (FC) of the membrane proteome of the WT compared to the Δ*R* strain. The FC corresponds to the WT/Δ*R* ratio. Blue and pink gradients indicate those proteins found significantly decreased or increased (FC ≥ |1|; *p-value* < 0.05), respectively, in the WT cells compared to Δ*R*. **B.** Distribution of cell lengths and cell curvature quantified from phase-contrast microscopy images of WT, Δ*R*, and Δ*R*Δ*ABT* strains at exponential phase in control conditions and exposed to 190 µM CuSO_4_ for 1 h. The solid line indicates the median, broken lines represent the 25^th^ and 75^th^ percentiles, respectively. **C.** SEM images of *C*. *vibrioides* WT and Δ*R* (left), Cryo-EM images of *C*. *vibrioides* WT and Δ*R* (centre), zoomed in images of Cryo-EM membrane defects of WT and Δ*R* (right). A total of 15 bacteria per strain were taken to measure OM-IM space, and the mean of 45 individual measurements all around the bacteria was taken. Scale bar, 100 nm. **D.** Fluorescence of propidium iodide and Syto9 in WT, Δ*R*, Δ*R* Δ*ABT,* Δ*R acrB\** and positive control WT permeabilized cells incubated with or without 190 µM CuSO_4_ for 1 h. All data are represented as the mean ± SD, with at least three biological replicates. *p* values were calculated using ANOVA combined with Tukey and Dunnett’s multiple comparison test (\*\*\*\**p* < 0.0001, \*\*\**p* < 0.001, \*\**p* < 0.01, \**p* < 0.05).

Scanning electron microscopy (SEM) imaging revealed prominent blebs on the surface of the Δ*R* cells under control conditions, but no major surface alterations, such as surface roughness, were observed compared to the WT strain (Fig. 3C). In line with these observations, a Δ*rsaA* mutant lacking the S-layer showed no increased Cu sensitivity (Fig. S2C). Furthermore, deletion of the *CCNA_00164* gene involved in *C. vibrioides* capsule biosynthesis had no impact on Cu resistance (Fig. S2C). Consistent with these observations, surface zeta potential measurements revealed no significant difference between WT, Δ*R*, and Δ*RΔABT* strains, with or without Cu treatment, indicating no major change in surface charge (Fig. S2D).

Cryo-electron microscopy (Cryo-EM) imaging uncovered more pronounced ultrastructural changes. WT cells displayed an intact cell envelope architecture with a consistent periplasmic space (IM-OM distance ∼31.5 nm +/- 1.2) (Fig. 3C), while Δ*R* cells exhibited a distorted, wave-like envelope architecture and a more variable intermembrane space (∼31.9 +/- 2.3 nm) (Fig. 3C), indicating a disruption of cell envelope integrity and potentially peptidoglycan defects. Under Cu stress, both WT and Δ*R* cells showed envelope defects, including slight undulations in the WT strain, prominent blebs in the Δ*R* mutant, and highly variable spacing in both strains (∼30.8 +/- 6.7 nm and ∼32.3 +/- 3.8 nm, respectively) (Fig. 3C and Fig. S2E).

We hypothesized that envelope defects in the Δ*R* mutant might increase membrane permeability and contribute to Cu sensitivity. Live/dead staining by flow cytometry revealed that the Δ*R* mutant shows a WT level of propidium iodide (PI) uptake under Cu stress, while a AcrAB_2_NodT impaired efflux mutant (Δ*R acrB*\*, see section below) displayed elevated PI uptake under both control and Cu stress conditions (Fig.3D). This data indicates that high AcrAB_2_NodT activity in the Δ*R* mutant triggers cell envelope defects with increased membrane permeabilization.

In summary, TipR loss-of-function leads to AcrAB_2_NodT upregulation, which disrupts bacterial envelope architecture and causes distinct morphological defects. The resulting Cu sensitivity is partially due to physiological stress from pump overactivity, rather than increased Cu accumulation.

### AcrB_2_ is a key player driving the collateral Cu sensitivity of the Δ*tipR* mutant

To determine whether the impact of AcrAB_2_NodT efflux pump on Cu sensitivity was due to an increased membrane crowding and/or to a depletion of bacterial energy sources, we sought to impair the AcrAB_2_NodT pump activity and to monitor the impact on Cu sensitivity. In *Salmonella typhimurium*, F615, F617, and R717 residues located in the AcrB porter domain lobe and switch loop connecting the proximal and distal binding pockets, respectively, were shown to play a key role in the AcrAB-TolC efflux regulation (15). A sequence alignment between *C. vibrioides* AcrB_2_ and *S. typhimurium* AcrB highlighted the conserved F610, F617, and R716 residues in AcrB_2_ (Fig. S3A), which are located near the predicted ligand in the drug-binding pocket. Based on our AlphaFold3 prediction, AcrB_2_ is likely to form a trimer, consistent with the oligomeric architecture of other RND efflux pumps (Fig. 4A and Fig.S3B). Structural modeling suggested that these substitutions increase the distance between the mutated residues and the ligand, potentially impairing pump function (Fig. 4A). Therefore, we mutated these residues into alanine in the Δ*R* genetic background, the resulting Δ*R acrB*\* mutant restored a WT growth profile under control and Cu stress conditions in both solid and liquid PYE medium (Fig. 4B–C). The Δ*R acrB*\* mutant also showed an increased sensitivity to CXM (one of the AcrAB_2_NodT substrates) compared to the Δ*R* strain in both solid and liquid media (Fig. 4B–C), confirming that AcrAB2NodT-mediated efflux is impaired by the F610A, F617A, and R716A point mutations.

**Figure 4.**
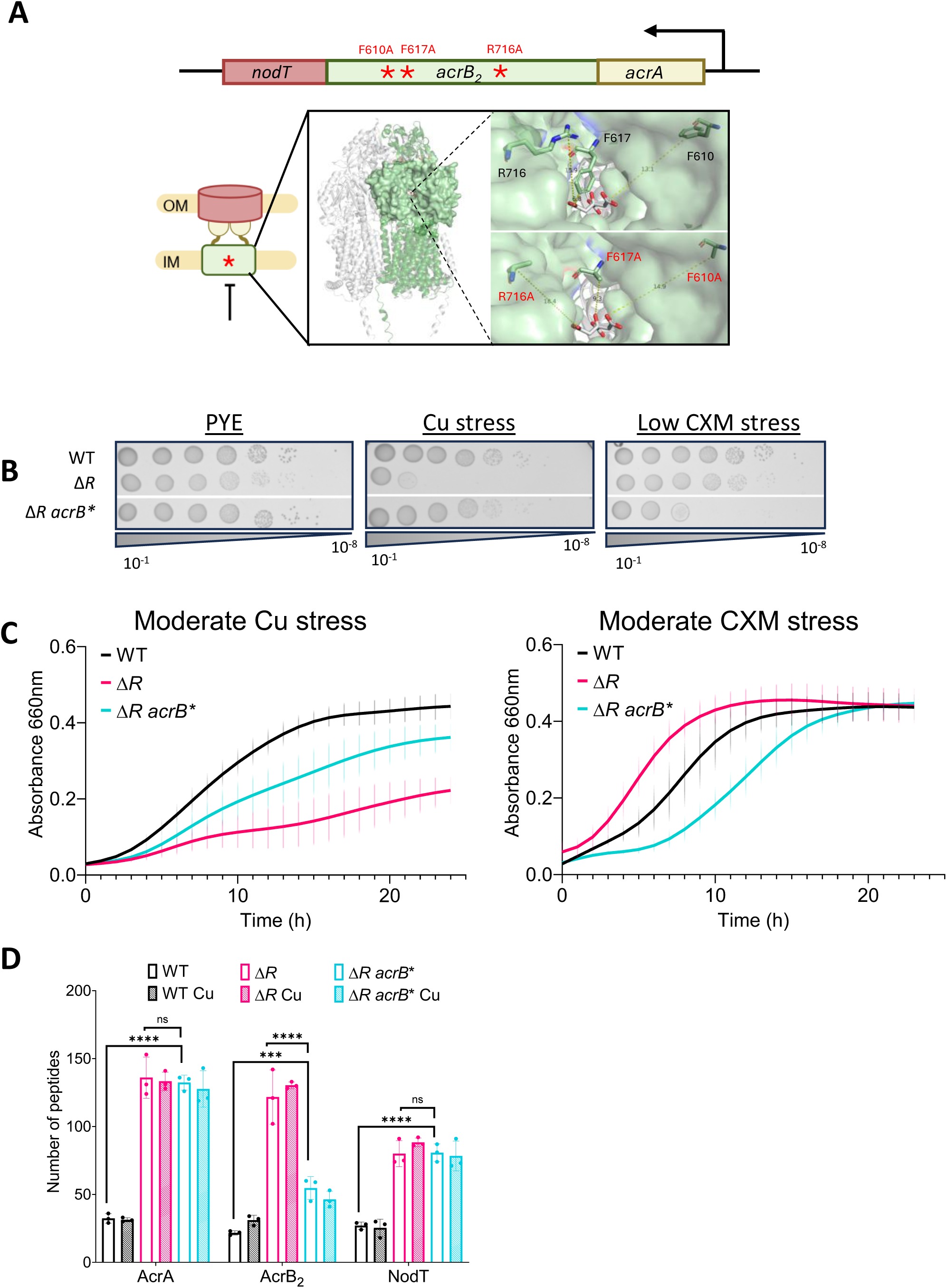
AcrB_2_ is the critical component linking efflux activity and Cu sensitivity. **A.** AlphaFold3 prediction of AcrB_2_ with and without point mutation (F610A, F617A, R716A) shown in red. **B.** Viability assay on PYE plates of WT, Δ*R*, and Δ*R acrB*\* strains, in control, 100 µM CuSO_4_ and 10 µg/ml CXM conditions. **C.** Growth profiles at an OD_660_ nm of WT, Δ*R*, and Δ*R acrB*\* strains in PYE medium supplemented with 150 µM CuSO_4_ and 35 µg/ml CXM, respectively. **D.** Normalized total spectrum count of AcrAB_2_NodT protein in PYE medium, in combination with a 1 h exposure to 190 µM CuSO_4_. All data are represented as the mean ± SD, with at least three biological replicates. *p* values were calculated using ANOVA combined with Tukey’s multiple comparison test (\*\*\*\**p* < 0.0001, \*\*\**p* < 0.001, \*\**p* < 0.01, \**p* < 0.05).

Interestingly, AcrB_2_ abundance was reduced in the Δ*R acrB*\* mutant relative to the Δ*R* mutant, although significantly higher than in the WT strain (Fig. 4D), indicating that the F610A, F617A, and R716A point mutations impact the AcrB_2_ stability. The higher sensitivity of the Δ*R acrB*\* mutant to CXM relative to the WT strain (Fig. 4B-C), despite a higher AcrAB_2_NodT abundance, suggests that F610A, F617A, and R716A point mutations also impair AcrB_2_ activity.

AcrA and NodT levels remained unchanged in the Δ*R acrB*\* mutant (Fig. 4D), suggesting that Cu sensitivity of the Δ*R* mutant mainly results from an overabundance of active AcrB_2_ in the IM, which could impair IM integrity and functions.

### Metabolic burden of AcrAB_2_NodT upregulation

RND efflux pumps use the PMF to actively expel toxic compounds. We hypothesized that overexpression of an active AcrAB_2_NodT efflux system in the Δ*R* mutant imposes a physiological burden through excessive consumption of the proton motive force (PMF), which could impair essential cellular processes such as ATP synthesis, motility, and nutrient transport and, in turn, affect Cu stress response.

To test this hypothesis, we measured the membrane potential as a PMF proxy using the ratiometric dye DiOC₂(3), where a decrease in red fluorescence indicates a loss in membrane potential. Under control conditions, the Δ*R* mutant exhibited a marked reduction in fluorescence compared to the WT strain, suggesting lower membrane potential likely due to PMF depletion (Fig. 5A). In contrast, both the Δ*R*Δ*ABT* and Δ*R acrB*\* mutants exhibited WT fluorescence, indicating that reduced AcrAB_2_NodT abundance and/or efflux activity restores membrane potential (Fig. 5A). Under Cu stress, both WT and Δ*R* strains showed a further drop in membrane potential, while the Δ*R*Δ*ABT* and Δ*R acrB*\* mutants remained largely unaffected (Fig. 5A). Consistent with a loss of PMF, the Δ*R* mutant exhibited a defective motility under both control and Cu stress conditions, while motility was restored to WT levels in the Δ*R*Δ*ABT* and Δ*R acrB*\* mutants (Fig. 5B). These results highlight that misregulated AcrAB_2_NodT activity is a major driver of PMF collapse, emphasizing the need for tight regulation of this efflux pump.

**Figure 5.**
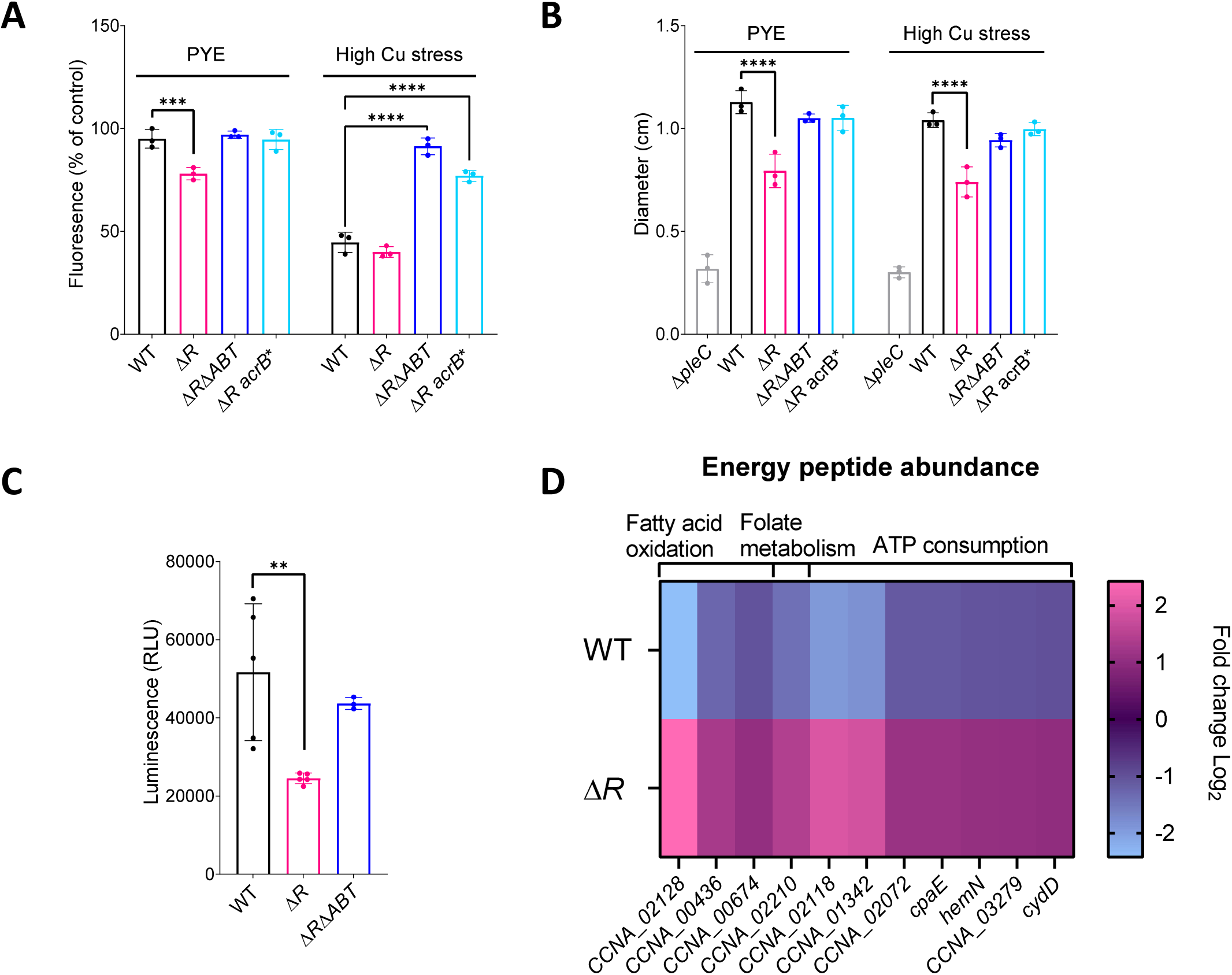
Cu sensitivity correlates with PMF disruption caused by AcrAB_2_NodT overactivity. **A.** Relative fluorescence intensity of the WT, Δ*R*, Δ*R*Δ*ABT*, and Δ*R acrB\** strains cells incubated with 29.7 µM DiOC _2_(3) dye under control conditions and exposed to 190 μM Cu. **B.** Motility assay of the WT, Δ*R*, Δ*R*Δ*ABT*, and Δ*R acrB\** strains after 72 h with and without Cu 100 µM. **C.** Heat map depicting the fold change (FC) of the proteome involved in energy metabolism of the WT compared to the Δ*R* strain. The FC corresponds to the WT/Δ*R* ratio. Blue and pink gradients indicate proteins found significantly decreased or increased (FC ≥ |1|; *p* < 0.05), respectively, in the WT cells compared to Δ*R*. **D.** Relative fluorescence intensity of the WT, Δ*R*, and Δ*R*Δ*ABT* strains incubated with ATP reagent HS under control conditions. All data are represented as the mean ± SD, with at least three biological replicates. *p* values were calculated using ANOVA combined with Tukey’s, Sidak’s, or Dunnett’s multiple comparison test (\*\*\*\**p* < 0.0001, \*\*\**p* < 0.001, \*\**p* < 0.01, \**p* < 0.05).

In line with the PMF impairment, the Δ*R* mutant exhibited a significant reduction in intracellular ATP levels relative to the WT strain (Fig. 5C). Accordingly, energy metabolism pathways are upregulated in the Δ*R* mutant, including ATP producing pathways such as fatty acid oxidation, folate metabolism, as well as ATP consuming processes such as DNA repair, ABC transporters, kinase activity, and motility, all of which rely on sustained energy availability (Fig. 5D).

These data collectively support a model in which loss of *tipR* leads to the upregulation of the AcrAB_2_NodT efflux system, resulting in depletion of the PMF and increased energy demand. This metabolic strain leads to ATP depletion, ultimately contributing to heightened sensitivity to Cu-induced stress.

## Discussion

RND efflux pumps have long been recognized as critical components in bacterial defense mechanisms against antimicrobial agents. Previous research has primarily focused on the protective benefits conferred by overexpression of such RND pumps under stressful conditions (16). However, the potential fitness costs associated with elevated expression of these large, energy-intensive complexes tend to be overlooked (10). In this study, we highlight the double-edged nature of efflux pump upregulation, demonstrating that while such systems can enhance stress resistance, it can also impose a fitness cost.

### OM remodeling is associated with AcrAB_2_NodT upregulation

While AcrAB_2_NodT is best known for exporting AB, our results show that its upregulation via *tipR* deletion sensitizes *Caulobacter* to Cu stress. This phenotype is not due to increased intracellular Cu or ROS, but instead reflects the physiological burden imposed by elevated efflux activity. Importantly, we found that AcrAB_2_NodT upregulation impacts outer membrane (OM) composition. Consistent with this, the Δ*R* mutant exhibited envelope defects, increased lipid metabolism, and upregulation of several TonB-dependent receptors (TBDR). In *C. vibrioides*, TBDR mediate nutrient uptake and are differentially regulated by AB and metals (17). Notably, BugA and ChvT influence sensitivity to Zn-binding AB, like bacitracin and vancomycin (18–20). A Zn-induced switch from ChvT to BugA has been shown, and we observed a similar upregulation of BugA under Cu stress (Fig. S4A), suggesting a shared regulatory mechanism (18). The UzcRS two-component system was previously shown to induce *acrAB_2_nodT* transcription under Zn stress, we observed no increase in AcrAB_2_NodT protein levels under Cu, suggesting regulation may be post-transcriptional or stress-specific (18). In parallel, AcrAB_2_NodT overexpression has been linked to OM remodeling through AbnZ, a small RNA derived from the 3′ end of the *acrAB_2_nodT* mRNA, which downregulates the TAM complex (21). This complex is essential for assembling autotransporters involved in nutrient uptake, adhesion, and virulence (22), and its disruption leads to membrane defects and reduced viability (21).

Together, these findings support a model in which AcrAB_2_NodT upregulation, combined with metal exposure drives envelope remodeling and disrupts membrane homeostasis. This structural and metabolic burden ultimately reduces the cell’s ability to tolerate metal toxicity.

### Energetic cost of efflux pump upregulation links PMF disruption to Cu sensitivity

AcrAB_2_NodT upregulation substantially depletes PMF, demonstrating that efflux pumps directly modulate cellular energetics. This effect is conserved across bacteria such as in *Salmonella typhimurium*, where reduced AcrAB activity increases membrane polarization (23). The resulting PMF disruption impairs ATP synthesis (24), creating a metabolic burden that could enhance metal sensitivity. Proteomics data revealed compensatory upregulation of energy-generating pathways (e.g., fatty acid oxidation, folate metabolism), likely to boost NADH/FADH₂ and restore electron transport chain (ETC) function. Comparable metabolic shifts occur in *Pseudomonas aeruginosa*, where efflux hyperactivity triggers anaerobic respiration to rebalance pH (10, 24). Although *C. vibrioides* is a strictly aerobic organism, AcrAB_2_NodT induction under microaerobic conditions (Fig. S5A) suggests a conserved adaptation to efflux-driven stress. Together, these findings position efflux pumps as key modulators of bioenergetics, with downstream consequences for metal resistance and metabolic adaptation.

Our findings provide new insight into the complex regulation of AcrAB_2_NodT, showing that its constitutive overexpression, while often protective, can impair bacterial fitness under metal stress. This fitness cost arises from membrane remodeling and metabolic adaptation, which shape Cu sensitivity and environmental resistance. This underscores the double-edged nature of efflux pumps and the critical need for tight regulation by factors like TipR and potentially UzcRS. Importantly, exploiting the trade-off between AB resistance and metal sensitivity could open new strategies to enhance AB efficacy via combination therapies.

## Materials and Methods

### Strains and plasmids

The *C. vibrioides NA1000* (WT) strain was grown at 30°C, under moderate shaking, in Peptone Yeast Extract (PYE) (0.2% bacto peptone, 0.1% yeast extract, 1 mM MgSO_4_, 0.5 mM CaCl_2_) medium, supplemented with cefuroxime (10 mg/ml), ciprofloxacin (1 mg/ml), kanamycin (50 mg/ml), and/or CuSO_4_.5H_2_O (Cu^2+^) when required. Exponentially growing cultures were used for all experiments. Plasmids were mobilized from a DH10B *E. coli* strain into *C. vibrioides* by triparental mating. Strains and plasmids are listed in Table S1. The strategies and the primers for their construction are available upon request.

### Growth curves

Bacterial cultures in the exponential growth phase (OD_660_ of 0.4 – 0.6) were diluted in PYE medium to a final OD_660_ of 0.05 and inoculated in 96-well plates with appropriate CuSO_4_ concentrations when required. Bacteria were then grown for 24 h at 30°C under continuous shaking in an Epoch 2 Microplate Spectrophotometer from BioTek and OD_660_ was measured every 10 min.

### Spot assays

Bacterial cultures in the exponential growth phase (OD_660_ of 0.4 – 0.6) were diluted in PYE medium to a final OD_660_ of 0.1. Ten-fold serial dilutions up to 10^-8^ (in PYE) were prepared in 96-well plates and drops of 5 μL of each dilution were spotted on PYE and PYE Cu plates using an automatic multichannel. Plates were incubated for 48 h at 30°C and pictures were taken with the Amersham Imager 600 (GE Healthcare LifeSciences).

### Atomic absorption spectrometry

15 mL cultures of *C. vibrioides* cells, were grown up to exponential phase (OD_660_ = 0.5). A 5 min or 1 hour treatment with 175 μM final CuSO_4_ concentration was applied and the cultures were then centrifuged at 8000 rpm for 10 min at 4°C. The cells were then fixed for 20 min on ice in 2% paraformaldehyde and then washed three times with an ice-cold wash buffer (10 mM Tris-HCl pH 6.8, 100 μM EDTA). For a total fraction, where the cytoplasm and periplasm are not separated, the pellet is resuspended in 2 ml of MilliQ H_2_O and then lysed with the cell disrupter (Cell Disruption System, One-shot Model, Constant) at 2.48 kPa. A final centrifugation at 10,000 rpm is performed for 10 min and 1.6 mL of the supernatant is mixed with 1 mL HNO_3_ 5% and 2.4 mL of MilliQ H_2_O for a final volume of 5 mL and a final concentration of 1M HNO_3_. Samples were finally analyzed by Shimadzu Atomic Absorption Spectrophotometer AA-7000F. The number of bacteria for each sample was calculated based on the OD_660_ measured prior to the lysis. The ratio between the OD_660_ and the number of bacteria was determined (25). Cellular metal concentrations were calculated using the following formula (25, 26):

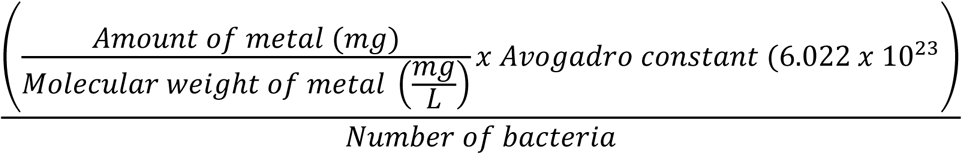

### Oxidative stress measurement with the CellROX Deep Red probe

An exponential phase culture of *C. vibrioides* was diluted to a final absorbance of OD_660_ of 0.2 and incubated for 30 min at 30°C under moderate shaking. We added 5 μM of CellROX Deep Red probe and appropriate concentrations of CuSO_4_ or H_2_O_2_. The bacteria were then washed three times with PBS to remove the noninternalized probe. The bacteria were finally transferred into a black bottom 96-well plate, and the fluorescence was monitored at emission/excitation of 640/690 nm with a SpectraMax iD3 (Molecular Devices). The relative fluorescence intensity was determined by defining 100% as the fluorescence from the control condition.

### Oxidative stress measurement with the rxYFP biosensor

An exponential phase culture of *C. vibrioides* culture overexpressing the rxYFP biosensor was diluted to a final absorbance of OD_660_ of 0.4 in a black bottom 96-well plate and mixed with different concentrations of CuSO_4_ or H_2_O_2_ (25). The rxYFP fluorescence was then monitored every minute for 20 min at emission/excitation of 510/556 nm with a SpectraMax iD3. The relative fluorescence intensity was determined by defining 100% as the initial measured fluorescence at time 0.

### Suppressor assay

The Δ*R* culture in the exponential growth phase (OD_660_ of 0.4 – 0.6) was diluted in PYE medium to a final OD_660_ of 0.1. Several drops of 5 μL were spotted PYE Cu 120 µM plates using an automatic multichannel. Plates were incubated for several days at 30°C until suppressors appeared. The colonies were streaked on PYE and PYE Cu plates to confirm the resistance. The suppressors that had Cu resistance were sent for whole-genome sequencing.

### Mass Spectrometry

Fifteen mL cultures of *C. vibrioides* cells were grown up to exponential phase (OD_660_ = 0.5). One hour treatment with 190 μM final CuSO_4_ concentration was applied, and the cultures were then centrifuged at 8,000 rpm for 10 min at 4°C. The cells were then fixed for 20 min on ice in 2% paraformaldehyde and then washed three times with an ice-cold wash buffer (10 mM Tris-HCl pH 6.8, 100 μM EDTA). For a total fraction, where the cytoplasm and periplasm are not separated, the pellet is resuspended in 2 mL of MilliQ H_2_O + protease inhibitor tablet and then lysed with the cell disrupter (Cell Disruption System, One-shot Model, Constant) at 2.48 kPa. The lysed cells were incubated with 1% SDS for 30 min on a spinning wheel. A final centrifugation at 10,000 rpm is performed for 10 min, and the protein concentration is determined by nanodrop and sent for LC/MS analysis.

### LC/MS

The samples were treated using the optimized filter-aided sample preparation protocol (27). Briefly, the samples were loaded onto Millipore Microcon 30 MRCFOR030 Ultracel PL-30filters that have been rinsed and washed beforehand with 1% formic acid (FA) and 8 M urea buffer (8 M urea in 0.1 M Tris buffer at pH 8.5), respectively. The proteins on the filter were then exposed to a reducing agent (dithiothreitol) and then alkylated with iodoacetamide. The proteins were then finally digested overnight with trypsin or trypsin/Glu-C (trypsin in 1/50 in ABC buffer; Glu-C in phosphate buffer 50 mM pH 7.4). The final step of the digestion is to transfer proteins in 20 μL of 2% acetonitrile (ACN) and 0.1% FA in an injection vial for reverse phase chromatography. The digest was analyzed using nano-LC–ESI–MS/MS tims-TOF Pro (Bruker, Billerica, MA, USA) coupled with an UHPLC nanoElute (Bruker). Peptides were separated on a 75 μm ID, 25 cm C18 column with integrated Captive Spray insert (Aurora, Ion Opticks, Melbourne) at a flow rate of 400 nL/min at 50°C. LC mobile phase A was water with 0.1% formic acid(v/v), and B was ACN with formic acid 0.1% (v/v). Samples were loaded directly on the analytical column at a constant pressure of 800 bars. The digest (1 μL) was injected, and the organic content of the mobile phase was increased linearly from 2% B to 15% in 22 min, 15% B to 35% in 38 min, and 35% B to 85% in 3 min. Data acquisition on the tims TOF Pro was performed using Hystar 5.1 and tims Control 2.0. tims TOF Pro data were acquired using 100 ms TIMS accumulation time and mobility (1/K0) range from 0.6 to1.6 Vs/cm². Mass-spectrometric analysis was carried out using the parallel accumulation serial fragmentation (PASEF) acquisition method (28). One MS spectrum was followed by 10 PASEF MSMS spectra per total cycle of 1.1 s. All MS/MS samples were analyzed using Mascot (Matrix Science, London, UK; version 2.8.1). Mascot was set up to search the *C. vibrioides* NA1000_190306 database from UniRef 100 and Contaminants_20190304 database assuming the digestion enzyme trypsin/GluC. Mascot was searched with a fragment ion mass tolerance of 0.050 Da and aparent ion tolerance of 15 PPM. Scaffold (version Scaffold_5.1.1, Proteome Software, Inc., Portland, OR) was used to validate MS/MS-based peptide and protein identifications. Peptide identifications were accepted if they could be established at greater than 97.0% probability to achieve an FDR less than 1.0% by the Percolator posterior error probability calculation (29). Protein identifications were accepted if they could be established at greater than 50.0% probability to achieve an FDR less than 1.0% and if they contain at least one identified peptide. Protein probabilities were assigned by the Protein Prophet algorithm (30). Proteins that contained similar peptides and could not be differentiated based on MS/MS analysis alone were grouped to satisfy the principles of parsimony. Proteins sharing significant peptide evidence were grouped into clusters.

### Microscopy

For live imaging, cells were grown to exponential phase (OD_660_ = 0.4) in PYE media. One hour treatment with 190 μM final CuSO_4_ concentration was applied. Cultures were then directly spotted on a microscope glass slide. Pictures were acquired with an Axio Observer (Zeiss) microscope equipped with an Orca-Flash 4.0 camera (Hamamatsu) and the Zen Pro 3.9 software (Zeiss). The images were processed and analyzed using ImageJ.

### SEM

Cells were post-fixed in 2% osmium tetroxide prepared in 0.1 M cacodylate buffer for 1 hour. Samples were subsequently washed three times (5 min each) in the same buffer and dehydrated through a graded acetone series (50%, 70%, 95%, and 100%), with 15 min at each step. Final drying was performed using critical point drying with a LEICA EM CPD030, which enabled the substitution of acetone with liquid CO₂, followed by vaporization at the critical point. Dried samples were then coated with a 20 nm platinum layer using a LEICA EM MED020 metallizer. Imaging was carried out using a QUANTA FEG 250 environmental scanning electron microscope (ESEM) (FEI, Thermo Fisher Scientific) equipped with a secondary electron detector (Everhart–Thornley type).

### Cryo-EM

QUANTIFOIL® R2/1 200 mesh Cu + C and lacey carbon grids were glow-discharged in an ELMO (CORDOUAN) at a vacuum of 2.1*10⁻¹ mbar and a voltage of 1.7 V for 35 sec. The grids were then used in a plunge-freezing procedure on the VITROBOT (ThermoFisher) with a double application of a 3 µL volume and the following parameters:

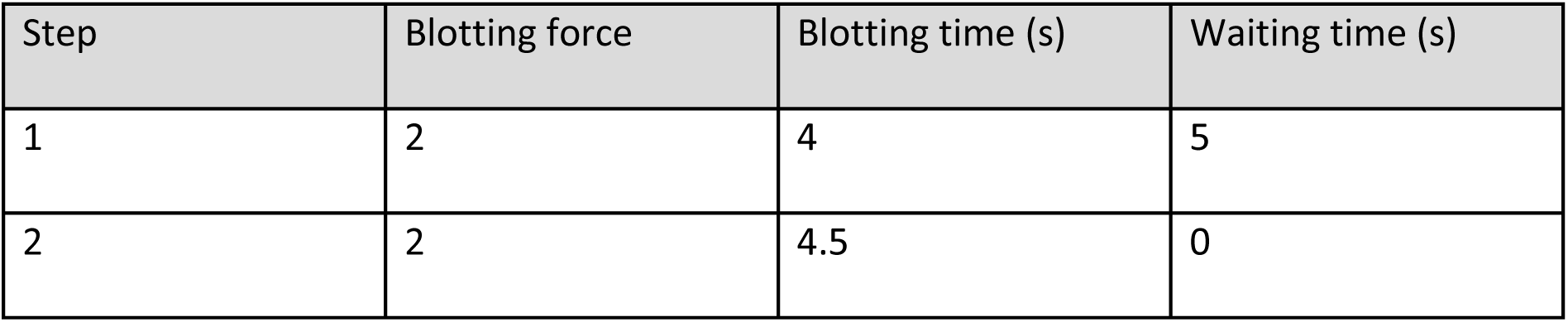

| Step | Blotting force | Blotting time (s) | Waiting time (s) |
| --- | --- | --- | --- |
| 1 | 2 | 4 | 5 |
| 2 | 2 | 4.5 | 0 |

The grids were then observed on the TALOS microscope (ThermoFisher) at 200 kV under cryogenic conditions. Images were acquired using EPU software and a Falcon III EC camera.

Images acquired at 36,000x magnification using the Falcon 3EC camera have a pixel size of 0.29 nm and a total electron dose of 20 e⁻/Å². The images were processed and analyzed using ImageJ.

### Zeta potential

Exponentially growing cells were incubated for 1 h with 190 µM of CuSO_4_. Bacteria were then centrifuged for 2 min at 12, 000 rpm. The pellet was washed three times with 1x PBS and resuspended in 1 ml distilled water. The Zeta (ζ) potential (millivolts, mV) was measured with a Nanopartica SZ-100 (Horiba Scientific) according to the manufacturer’s instructions.

### Bacterial live/dead assay

Exponentially growing cells were incubated for 1 h with 190 µM of CuSO_4_. Bacteria were then centrifuged for 10 min at 10,000g. The pellets were resuspended in 0.85% NaCl or 70% isopropyl alcohol and incubated for 1 h with shaking every 15 min. Samples were centrifuged and resuspended twice more. The optical density at 670 nm was determined. Cells were normalized to have 1 x 10^6^ bacteria/mL and stained following the LIVE/DEAD BacLight bacteria viability kit. The percentage of live or dead bacteria was determined by BD FACS verse cell analyser.

### Membrane potential measurement using DiOC_2_(3)

An exponential phase culture of *C. vibrioides* was diluted to a final absorbance of OD_660_ of 0.2 and incubated for 30 min at 30°C under moderate shaking with appropriate concentrations of CuSO_4_. The bacteria were then washed three times. 3 mM of DiOC_2_(3) probe or 500 µM CCCP was used according to the manufacturer’s protocol (Bac*<u>Light</u>* bacterial membrane potential kit). The bacteria were finally transferred into a black bottom 96-well plate, and the fluorescence was monitored at emission/excitation of 500/690 nm with a SpectraMax iD3 (Molecular Devices). The relative fluorescence intensity was determined by defining 100% as the fluorescence from the control condition.

### ATP Kit

Exponentially growing cells were diluted to an OD_660_ of 0.1 in mineral water. Bacteria were transferred into a white bottom 96-well plate. Cells were treated using the intracellular ATP kit HS according to the manufacturer’s protocol. The luminescence was measured with a SpectraMax iD3 (Molecular Devices).

### Motility assay

*C. vibrioides* cultures were grown in PYE overnight at 30°C with 150 rpm. The cultures were inoculated with sterile toothpicks on a PYE swarming plate (PYE + 0.25% agar). The plates were incubated for 3 days at 30°C and then imaged using Amersham Imager 600.

### Statistical analysis

Statistical analysis were performed when required. The data were analyzed with ANOVA one- way combined with Dunnett’s, Tukey’s, and Sidak’s multiple comparisons test. A p-value below 0.05, 0.01, 0.001 and 0.0001 are represented by *, **, ***, and **** respectively. A T-test was performed when needed.

### *In silico* analysis of RND efflux pump homologs

A blast search of each protein for both AcrABTolC pump and CusCBA of *E. coli* was achieved by using biocyc “blastp” and searched for orthologs in *C. vibrioides NA1000*. Several features such as query coverage, bit score, E. value, identity, similarities, and gaps were considered to identify the best homologs. Once the homologs of these pumps were identified in *C. vibrioides NA1000* a reverse best match was performed using “blastp” in *E. coli*.

### Sequence-based bioinformatic predictions

A multiple sequence alignment of AcrB_2_ protein from *Caulobacter vibrioides* (Uniprot ID: B8H1X6) and AcrB protein from *Salmonella typhimurium* (Uniprot ID: Q8ZRA7) was first performed using the online MAFFT tool (https://www.ebi.ac.uk/jdispatcher/msa). The results were visualized using Jalview.

## Acknowledgements

We thank Laurelenn Hennaux (UCPTS laboratory, UNamur) for her help with the DLS for zeta potential measurements. We thank Céline Legrand (MORPH-IM, Unamur) for her support with the flow cytometer. We acknowledge Christine-Jacobs Wagner, Marianne Ilbert, Patrick Viollier, Jordan Costafrolaz and the URBM members for fruitful discussions and precious tips for different experiments.

## Author contributions

M.O. and J.-Y.M. conceptualization; M.O. methodology; M.O. validation; M.O., L.L., E.H., D.J., M.D. and J.-Y.M. formal analysis; M.O., L.L., E.H., D.J., and M.D. investigation; M.O. writing– original draft; M.O. visualization; J.-Y.M. writing–review & editing; J.-Y.M. supervision; J.-Y.M. funding acquisition.

---

**Figure S1: Cu sensitivity of the Δ*R* mutant is not Cu-specific. A.** Growth profiles at an OD_660_ nm of WT, Δ*R*, and Δ*R+R* strains in control PYE medium and supplemented with 175 µM and 190 µM CuSO_4_. **B.** Growth profiles at an OD_660_ nm of WT, Δ*R*, and Δ*R+R* strains in control PYE medium supplemented with 75 µM of ZnSO_4_. **C.** Viability assay on PYE plates of WT, Δ*R*, and Δ*R+R* strains, in 50 µM ZnSO_4_, 200 µM NiSO_4_ and 2 µM CdSO_4_ conditions. **D.** Number of Cu atoms per cell exposed to 175 µM CuSO_4_ excess for 5 min. All data are represented as the mean ± SD, with at least three biological replicates.

**Figure S2: Microscopic and genetic analysis of cell surface alterations and their Impact on Cu sensitivity. A.** Suppressors of the Δ*R* mutant were found on PYE plates supplemented with 120 µM CuSO_4_. In total, 9 suppressors were isolated, which to some extent showed improved growth on Cu compared to the Δ*R* mutant. **B.** Phase-contrast microscopy images of WT, Δ*R*, and Δ*R*Δ*ABT* strains at exponential phase in control condition and exposed for 1 h to 190 µM CuSO_4_ (bar = 5 μm). **C**. Growth profiles at an OD_660_ nm of WT, Δ*R*, Δ*rsaA*, and Δ*CCNA_00164* strains PYE medium supplemented with 150 µM CuSO_4_. All data are represented as the mean ± SD, with at least three biological replicates. **D.** Zeta potential measurements of WT, Δ*R*, and Δ*R*Δ*ABT* cells with or without 190 µM CuSO_4_ treatment. Mean ± SD, at least two biological replicates. **E.** Diameter of the distance between OM-IM space from Cyro-EM images. Distribution of OM-IM diameter space quantified from Cryo-EM microscopy images of WT and Δ*R* strains at exponential phase in control condition and exposed for 1 h to 190 µM CuSO_4_ . The solid line indicates the median, broken lines represent the 25^th^ and 75^th^ percentiles, respectively.

**Figure S3: Multiple sequence alignment of AcrB_2_ and AlphaFold3 prediction. A.** Multiple MAFFT alignment of AcrB_Cv_ with AcrB homologs from *Salmonella typhimurium*. Identical residues are highlighted in blue, and the three conserved amino acids of interest, F610, F617, and R716, are highlighted in red. **B.** Graphic plot of the prediction of AcrB_cv_ and AcrB_cv_* accounting for the pLDDT and PAE (source AlphaFold3).

**Figure S4: Protein abundance of BugA. A.** Normalized total spectrum count of BugA protein in PYE medium, in combination with a 1 h exposure to 190 µM CuSO_4_. All data are represented as the mean ± SD, with at least three biological replicates. *p* values were calculated using ANOVA combined with Tukey’s multiple comparison test (\*\*\*\**p* < 0.0001, \*\*\**p* < 0.001, \*\**p* < 0.01, \**p* < 0.05).

**Figure S5: Metabolism switch upon AcrAB_2_NodT expression. A.** Normalized total spectrum count of AcrAB_2_NodT protein in PYE medium, in aerobic and microaerobic conditions. All data are represented as the mean ± SD, with at least three biological replicates. *p* values were calculated using ANOVA combined with Sidak’s multiple comparison test (\*\*\*\**p* < 0.0001, \*\*\**p* < 0.001, \*\**p* < 0.01, \**p* < 0.05).

**Table S1.** Bacterial strains and plasmids used in this study.

| Strain | Genotype | Reference |
| --- | --- | --- |
| WT | *Wild-type NA1000, synchronizable variant of CB15* | (31) |
| $\Delta R$ | *C. vibrioides strain deleted for tipR (CCNA_00852)* | This study |
| $\Delta R+R$ | *C. vibrioides $\Delta R$ strain harbouring tipR (CCNA_00852) on pMR10 plasmid* | This study |
| $\Delta ABT$ | *C. vibrioides strain deleted for acrAB<sub>2</sub>nodT (CCNA_00849:51)* | This study |
| $\Delta R\Delta ABT$ | *C. vibrioides strain deleted for tipR and acrAB<sub>2</sub>nodT* | This study |
| WT+ABT | *C. vibrioides WT strain harbouring acrAB<sub>2</sub>nodT on pMR10 plasmid* | This study |
| $\Delta R$ acrB* | *C. vibrioides $\Delta R$ strain harboring the acrB gene (CCNA_00850) mutated for F610A F617A R716A* | This study |
| WT rxYFP | *C. vibrioides WT harboring the plasmid pJS14 rxYFP* | This study |
| $\Delta R$ rxYFP | *C. vibrioides $\Delta R$ harboring the plasmid pJS14 rxYFP* | This study |
| **Plasmids** |  |  |
| pNPTs138 | *mobRP4+ ori-R6K sacB; integrative vector in C. vibrioides for in-frame deletions; KanR* |  |
| pMR10 | *Low copy and replicative vector in C. vibrioides; KanR* |  |
| pJS14 | *pJS14 medium copy number replicative cloning vector in E. coli and C. vibrioides; CmR* |  |

